# Assessment of urinary exosome NHE3 as a biomarker of acute kidney injury

**DOI:** 10.1101/2022.06.12.495794

**Authors:** Yanting Yu, Zhiyun Ren, Anni Xie, Yutao Jia, Ying Xue, Ping Wang, Daxi Ji, Xiaoyan Wang

## Abstract

The diagnosis of acute kidney injury (AKI) traditionally depends on the serum creatinine (SCr) and urine output, which lack sufficient sensitivity and specificity. Urinary exosome as a biomarker has unique advantages. To assessed whether urinary exosome Na+/H+ exchanger isoform 3 (NHE3) protein as a nonivasive diagnostic biomarker of AKI, we constructed 4 AKI rat models: cisplatin (7.5mg/kg) injection intraperitoneally (IP), furosemide (20mg/kg, IP) with low NaCl (0.03%) diet, low NaCl (0.03%) diet with candesartan (1mg/kg, IP) and bilaterally ischemia and reperfusion(I/R) injury for 40 minutes. Besides, we assessed 6 sepsis associated AKI patients and 6 healthy volunteers. Urinary exosomes were extracted by ultra-centrifugation and NHE3 protein abundance was tested by immunoblotting in all AKI rats and human subjects. The isolated cup-shape particles with an average diameter of 70nm and enrichment in CD63 were identified as exosomes. NHE3 abundance was 6 times higher in exosome than in original urine. In cisplatin induced AKI rats, urinary exosome NHE3 was increased at day 2, 1 day earlier than the increases of serum creatinine creatinine (SCr) and blood urea nitrogen (BUN). In additional rats, urinary exosome NHE3 decreased along with the decline of SCr after EPO pretreatment. In volume depletion AKI induced by furosemide injection with low NaCl diet, urinary exosome NHE3 expression was higher than control. In low NaCl diet with candesartan related AKI, urinary exosome NHE3 was elevated at day 5, 2 days earlier than SCr. In I/R injury AKI, urinary exosome NHE3 was also raised compared with control. In humans, the urinary exosome NHE3 level was also elevated in sepsis associated AKI patients in comparison with the healthy volunteers. Then urinary exosome NHE3 protein may be used as a noninvasive diagnostic biomarker of AKI.

**Impact Statement:** The high non-recognition acute kidney injury (AKI) is due to lacking of alarming symptoms or specific early biomarkers. Urinary exosome as a biomarker has unique advantages. In our study, we detected urinary exosome NHE3 protein abundance in 4 different cause of AKI rat model. Urinary exosome NHE3 was increased in all 4 AKI, and even elevated earlier than SCr in some cases. Another novel point was we established a new AKI model of low NaCl diet with candesartan injection, which was common in patients with hypertension or proteinuria clinically. The detailed methods and mechanisms of this new AKI model were presented in another article being submitted. Third, we are not limited to animals, but also selected sepsis associated AKI patients to study. The conclusion that urinary exosome NHE3 may be used as a diagnosis biomarker of AKI has important clinical application value.

## Introduction

Acute kidney injury (AKI) is a global public health concern associated with high morbidity^1^and mortality^2^. Non-recognized AKI delays the diagnosis. A survey of 2.2 million patients shows that 74.2% of the patients with identifiable AKI are not recognized by physicians, and 17.6% recognized AKI are given a delayed diagnosis^3^. The high non-recognition is due to lacking of alarming symptoms or specific early biomarkers. AKI is defined by a fixed percentage of increase in serum creatinine (SCr), decrease in urine output, or both within 7 days, according to the Kidney Disease: Improving Global Outcomes (KDIGO) criteria^4^. SCr has poor sensitivity and specificity to detect or grade the severity of AKI, and lags behind both renal injury and renal recovery. Furthermore, urine output is difficult to quantify precisely if without an indwelling catheter. Hence, novel serum or urinary biomarkers for AKI are in need and have been a hot topic in the field.

Several urinary biomarkers have been discovered as non-invasive indicators for prediction and diagnosis of AKI including Kidney injury molecule-1, N-acetyl-beta-D-glucosaminidase^5, 6^, neutrophil gelatinase-associated lipocalin^7^ and liver fatty acid-binding protein^8^. The urinary product of tissue inhibitor metalloproteinase and insulin growth factor binding protein-7^9^ and urinary thioredoxin^10^ are also clinically available. But each biomarker currently has limitations. Moreover, their adoption into routine clinical care has been slow probably due to the cost, availability of testing platforms, variability in assay techniques and results, and lack of governance approval. In order to improve early detection of AKI, additional urinary biomarkers remain to be identified.

Urinary exosomes containing apical membrane and intracellular fluid are normally secreted into the urine from all nephron segments, and may carry biomarkers, such as proteins and microRNA, of renal dysfunction and structural injury. The application of urinary exosomes as markers has been benefited from the extraction of exosomes from healthy human urine by Pisitkun Trairak and the colleagues^11^. They first extracted urinal exosomes and identified 295 unique proteins, including several transporters and channels associated mainly with the apical membrane. The Na^+^/H^+^ exchanger isoform 3 (NHE3) is expressed in the apical membrane of the mammalian proximal convoluted tubule and thick ascending limb^12^. NHE3 belongs to the mammalian NHE protein family and catalyzes the electroneutral exchange of extracellular sodium for intracellular proton across cellular membranes. Actually, all the major sodium transporters expressed along the nephron including NHE3, Na-K-2Cl cotransporter, and the thiazide-sensitive Na-Cl cotransporter are detectible in urine of rats by means of antipeptide antibodies, suggesting that profiling sodium transporters in urine may become a useful test tool to detect and classify kidney diseases^13^. The detection of urinary exosome proteins^14^ and microRNAs^15^ in kidney disease allows experts to explore the possibility of exosomes as specific biomarkers for AKI since they are actively secreted by live cells. This study detected the urinary exosome NHE3 protein in various AKI rats and sepsis associated AKI patients. In this study, we detected the urinary exosome NHE3 protein in various AKI rats and sepsis associated AKI patients, aiming to assess it as a new early biomarker for AKI.

## Materials and methods

### Urine collection and exosome extract

24hr urine from rats and morning urine from patients were centrifugated at 4000g for 10 min to remove debris and stored in −80°C refrigerator. Exosome was isolated by ultra-centrifugation according to manufacturer’s instructions^11^. Urine samples were centrifugated at 17000g for 10 min at 4□. The supernatant 1 was collected, and the precipitation added 2ml isolation buffer containing 200mg/ml Dithiothreitol (DTT) was centrifuged at 17000g for 10 min again, then the supernatant 2 was collected and mixed with supernatant 1. The supernatant 1 and 2 were isolated by centrifugation 200,000g for 1 hr. The pellets were resuspended in 60ul of isolation solution containing 1g/mL leupeptin and 0.1mg/mL Phenylmethylsulfonyl fluoride(PMSF). Isolated exosomes were kept at 2-8°C for next serious experiments.

### Animals and Experimental Protocol

#### Animal

Eight-week-old male Sprague-Dawley (SD) rats weighing 300–350g were purchased (Beijing, China) and were used for various studies. The animal experiments were approved by the Committee on Animal Care of Nanjing Medical University and were conducted according to the National Institutes of Health Guidelines for the Care and Use of Laboratory Animals. All studies involving animals are reported in accordance with the Animal Research: Reporting of In Vivo Experiments (ARRIVE)guidelines. The animals were maintained in a 12–h light/dark cycle in a temperature and humidity-controlled acility and were given a minimum 7-day acclimation. The rats were placed in individual metabolic cages (Shanghai Yuyan Instruments Co.,Ltd), allowing daily ration feeding and collection of urine. Blood and tissue samples were harvested at the end time and processed for various studies.

#### Cisplatin-induced AKI

12 SD rats were divided into control group or cisplatin group. Cisplatin (Cis, 7.5mg/kg; Sigma) was single intraperitoneally injected as the previous research^16^, and the Cis group rats were fed for 1 day, 2 days or 3 days, respectively. The control rats received the vehicle, 0.9% saline and sacrificed at day 3 (n=3 per group). For the rescue experiment, 13 SD rats were random assigned into the control(n=4), cisplatin (Cis, n=4) and cisplatin with erythropoietin (Cis+EPO, n=5) group. EPO (5000U/kg; EPIAO) was administered 2 times, 15 min before cisplatin and 2 days after cisplatin administration. All rats were sacrificed at day 3.

#### Volume depletion induced AKI

8 SD Rats were divided randomly into control group or Volume depletion (VD) group according to the previous method^17^. The VD group rats were fed food with 0.4% NaCl (Jiangsu Xie Tong Pharmaceutical Bioengineering CO., LTD) for18 hours (−18hr) and then were injected intraperitoneally furosemide (BP547, sigma, 20mg/kg) once (0hr), 8 hours later(8hr) received the second furosemide injection at same dose, and were sacrificed at 30 hr. During all the 48hr the rat were fed with food containing 0.4% NaCl. The control rats were administrated low salt (0.03%NaCl) intake and the other procedure was the same. Blood was collected at 30^th^ hr. Urine samples at control group were collected for all the 48 hrs. Urine from VD group were collected from different period: −18hr to 0hr, 0-8hr, 8-24hr and 24–30hr.

#### Low NaCl diet with candesartan related AKI

All 12 SD rats were randomly divided into 2 group. Low NaCl group rats were fed food with 0.03% NaCl. Normal salt group were fed food with 0.4%NaCl. All rats were intraperitoneally injected with candesartan (PHR1854, sigma,1mg/kg/day) for one week. Rats were sacrificed at day 7. The low NaCl diet with candesartan model was established previously to study epithelial sodium channel expression in kidney^18^.

#### Ischemia/reperfusion related AKI

SD rats were assigned randomly to 2 groups: sham-operation group (Control) and I/R group(n=6/group). Rats were under surgical anesthesia with 0.5mg/kg chloral hydrate (Macklin, Shanghai, China). The abdomen was opened and the left renal pedicles were found to clamp, and then the right, each clamped for 40 min at 37°C^17^. After releasing the cross-clamps, the kidneys were re-perfused and the color returned to original. The abdomen was closed in two layers. Sham-operated group were subjected to similar surgical procedure except clamp.

#### AKI patients

A total of 12 subjects were recruited into the study from Nanjing BENQ hospital. 6 was diagnosed AKI based on the criteria of KDIGO. The etiology of AKI was clinically considered to be sepsis and no nephrotoxic drugs administration. 6 healthy control participants were from Health Management Center. For all subjects the morning urine was collected.

### Transmission electron microscopy

The 200,000g pellet isolated from the urine were rinsed and post-fixed in 1% osmium tetroxide. Samples were embedded in 10% gelatin, fixed and cut into several blocks (<1 mm^3^). The samples were dehydrated in increasing concentrations of alcohol and infiltrated with increasing concentrations of Quetol-812 epoxy resin mixed with propylene oxide. Samples were embedded in pure fresh Quetol-812 epoxy resin and polymerized. Ultrathin sections (100 nm) were cut by using a Leica UC6 ultramicrotome and post-stained with uranyl acetate for 10 min and lead citrate for 5 min at room temperature, followed by observation with a transmission electron microscope (HT-7700, Hitachi, LTD Japan) operated at 100 kV.

### Nanoparticle tracking analysis

The 200,000g pellets were sized and enumerated by nanoparticle tracking analysis (NTA) using Nano FCM (N30E) instrument. NTA is a light-scattering technique that uses video analysis for the sizing and enumeration of extracellular vesicles. Urine samples were collected and diluted in PBS to a particle concentration within the range of 2–10×10^8^/mL (optimal working range of the system). An approximately 100 μl diluted sample was loaded into the sample chamber, and 60 s videos were recorded for each sample with a shutter speed of approximately 30 ms and a camera gain between 250 and 650. The settings for software analysis were as follows: detection threshold, 30–50; blur, 535; and minimum expected particle size, auto. The size distributions are presented as 5–6 video recordings per sample.

### Biochemical analyses

For rats, the SCr was measured with the QuantiChrom Creatinine Assay Kit (cat.: DICT-500, Bioassay Systems, Hayward, CA), and BUN was measured with the QuantiChrom Urea Assay Kit (cat.: DIUR-500, Bioassay Systems) according to the manufacturer’s instructions. The patient’s serum creatinine and BUN were tested in Clinical Laboratory using Roche instrument(cobas8000).

### Western blotting

The different fraction of urine from rats separated by differential centrifugation were mixed with 1:4 vol/vol of Laemmli buffer and heated to 60°C for 15 minutes. The loading volume for immunoblotting was normalized to urine creatinine content. Kidney homogenates were prepared on ice using RIPA lysis buffer. The total protein concentration was measured with BCA kit (P0009, Beyotime). Equal amounts of protein from each sample were electrophoresed on SDS–polyacrylamide gel and electro-transferred onto nitrocellulose membranes which were incubated with primary antibodies anti-CD63(25682-1-AP, Proteinch), anti-NHE3(sc58636, Santa Cruz), anti-β-actin (A2066, Sigma). The chemoluminal signals from HRP-conjugated secondary antibodies (1:10000, Santa Cruz, CA) were exposed onto x-ray film. The band densities of the proteins were quantified using Image J software (NIH, Bethesda, MD, USA) and expressed as a percentage of the relevant control density.

### Histology examination

Renal tissues were fixed in 10% paraformaldehyde and embedded in paraffin, and 3μm-thick sections were prepared. The sections were stained with periodic acid-Schiff (PAS). The tubular luminal dilation, loss of brush border and flattened tubular epithelial cells and capillary loop shrinkage were viewed with Nikon Eclipse 80i microscope equipped with a digital camera (DS-Ri1, Nikon, Shanghai, China).

### Immunofluorescent staining

Kidney cryosections at 3μm thickness were fixed for 15min in 10% paraformaldehyde, followed by permeabilization with 0.2% Triton X-100 in phosphate-buffered saline (PBS) for 5min at room temperature. After blocking with 2% donkey serum for 60min, the slides were immunostained with anti-NHE3 (sc58636, Santa Cruz), subsequently with the appropriate fluorophore-conjugated secondary antibodies (1:500; Molecular Probes, Grand Island, NY, USA). The slide was mounted using ProLong Gold Antifade Mountant (Thermo Fisher Scientific) and imaged with LSM 800 confocal microscope (Carl Zeiss,Oberkochen, Germany).

### Statistical analyses

All quantitative data are expressed as mean ± SEM. Statistical analysis of the data was performed using SPSS 22.0 software (SPSS, Chicago, IL, USA). Comparison between groups was made using one-way ANOVA, followed by post hoc SNK or post hoc LSD. For comparison between two groups, Student’s t test was used, and paired t test was used for group comparison. P values of less than 0.05 were considered significant.

## Results

### Isolation and identification of urinary exosome and urinary NHE3 protein

We isolated exosomes by 2 steps of 17,000g and 200,000g differential centrifugation from urines of normal SD rats. The 200,000g pellets were tested by transmission electron microscopy (TEM) and nano tracking analyzer (NTA). The exosomes in this fraction were characterized by the following points: The TEM showed the round, cup-shaped vesicles, which diameter wasless than 100nm (Figure 1(a)); NTA revealed the particle diameter ranged at 50–100 nm with a mean around 75 nm (Figure 1(b)); The urine different fractions were tested by western blot for exosome marker CD63. CD63 protein was rich in abundant at 200,000g pellet (Figure 1(c)). Immunofluorescent staining for NHE3 (apical membrane) confirmed the NHE3 location in the kidney, probably the proximal tubules (Figure 1(d)).

**Figure 1.**
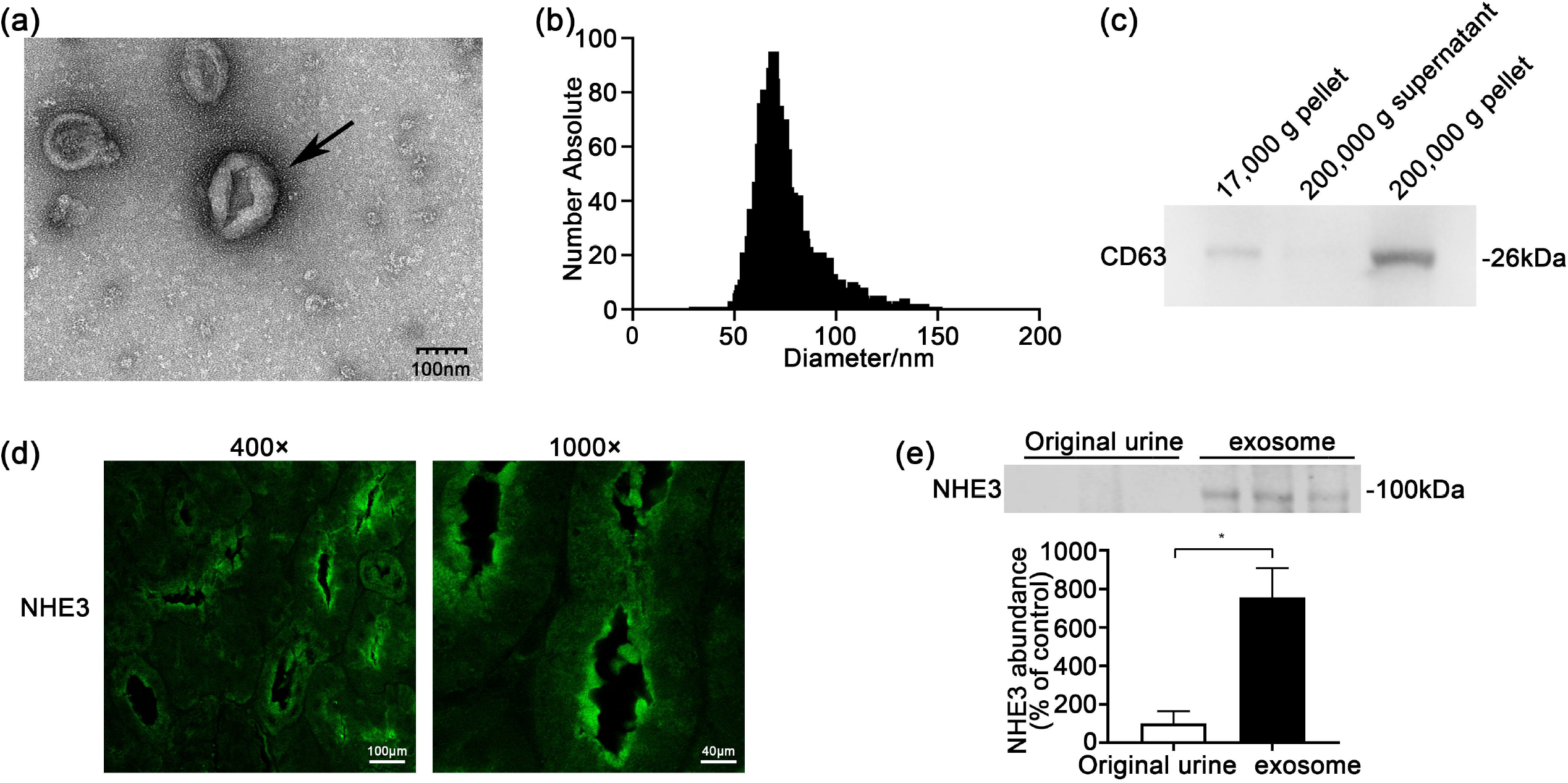
Characterization of urinary exosomes and NHE3 protein expression in kidney and urine. (a) Electron micrograph image of urine exosomes. The urine was centrifuged at 17,000×g for 10 min and the supernatants was ultracentrifuged 200,000×g for 1 hour to pellet the exosomes. The image shows round, cup-shaped vesicles. Scale bar = 100 nm. (b) Particle size distribution in purified pellets consistent with size range of exosomes (average size around 75 nm), measured by Zeta View® particle tracking analyzer. (c) Detection of CD63 protein in different urinary fractions from the same urine sample by Western blot analysis in Sprague–Dawley (SD) rats. (d) Immunofluorescent staining images for NHE3 in the kidney of normal SD rats. (e) NHE3 protein expression at whole urine and urine exosomes in normal SD rats. The loading amount was corrected according to the urinary creatinine content, and the same amount of creatinine was loaded. \**P* < 0.05.

Then, we compared the NHE3 level at original urine and urinary exosomes. NHE3 abundance in the exosome was 7 times more than that in original urine(755.6±152.6 vs. 100±63.5, *P*<0.05) (Figure 1(e)).

### Urinary exosome NHE3 expression in Cisplatin-induced AKI

The SCr(379.7±32.9 vs. 28.7±1.8 umol/l, *P*<0.01) and blood urea nitrogen (BUN) (53.9± 1.9 vs. 5.0±0.4mmol/l, *P*<0.01) were elevated at day 3 after cisplatin injected indicated AKI occurred (Figure 2(a) and 2(b)). The 24 hr urine output was increased at cisplatin group, whether compared with control or day 0, which was consistent with previous study^19^ (Figure 2(c)). The urinary exosome NHE3 was incremental from day 2(15880±2866vs.100±44.3, *P*<0.01), that earlier than serum creatinine increasing at day 3, and it was still increased at day3(29688±475 vs. 100±44.3, *P*<0.01). Next, we constructed an AKI recovery model by pretreated with EPO 15min before cisplatin injection, and 2 days after cisplatin injection. After EPO treatment, the increase of serum creatinine(78.2±35.4 vs.175±36.3 umol/l, *P*<0.05) and BUN(16.2±7.8 vs. 35.9±6.3, *P*<0.05) were reversed (Figure 2(e) and 2(f)), and the expansion of renal tubules and epithelial cell loss were alleviated (Figur 2(h)). Consistent with this, urinary exosome NHE3 was increased in cisplatin group(4517±1758 vs. 100± 41.63, *P*<0.05), but decreased after EPO pretreatment(501.2±253.1vs. 4517±1758, *P*<0.05), and had no statistic difference compare with control group. (Figure 2(g)). But the kidney NHE3 protein change was not inconsistent (Figure 2(i)).

**Figure 2.**
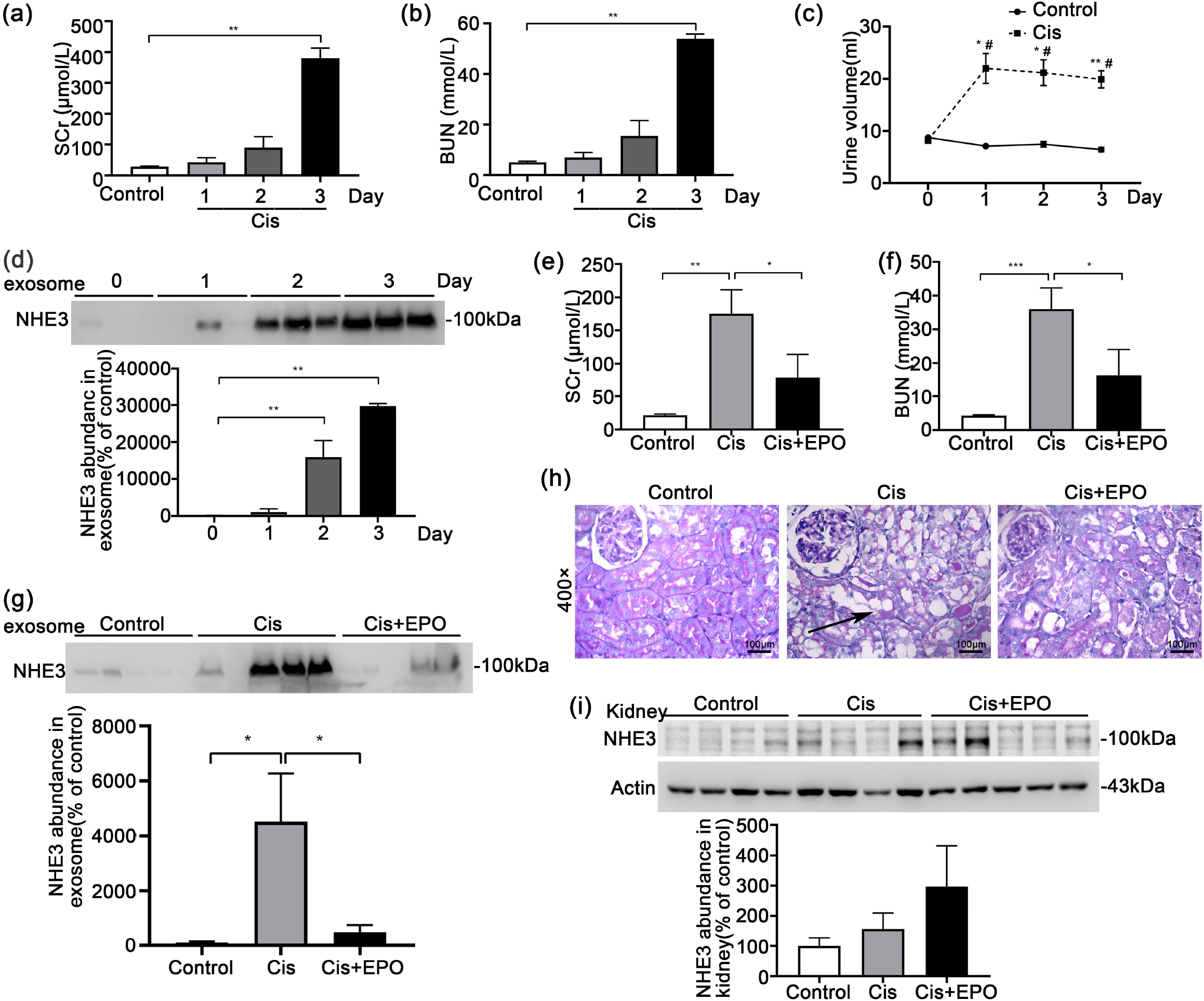
NHE3 expression in the cisplatin-induced acute kidney injury rats. (a, b) Comparison of SCr and BUN between the control and cisplatin-treated rats. 12 rats were intraperitoneally injected with saline (control) or cisplatin (7.5mg/kg; Sigma) which were sacrificed at 1 day, 2 days or 3 days, respectively(n=3 per group). (c) The urine output daily in control and cisplatin rats. * vs. control group. # vs. day 0. (d) Urinary exosome NHE3 protein expression at control rats and cisplatin treated rats at day1, day2 and day3. (e, f) Comparison of serum creatine and BUN levels among the control(n=4), cisplatin (Cis, n=4) and cisplatin+EPO (Cis+EPO, n=5) group. Cisplatin (7.5mg/kg) was given at day 1 i.p. EPO (5000U/kg; EPIAO) was administered 2 times, 15 min before cisplatin administration and 2 days after cisplatin administration. All rats were sacrificed at day 3. (g) Urinary exosome NHE3 protein expression by immunoblot among the three groups. (h) Renal histological changes in the different groups. Kidney sections were stained with PAS to assess morphological changes. Arrows indicated renal tubular dilatation and tubular epithelial cells loss. (i) NHE3 protein level in kidney by immunoblot among the three groups. The data are presented as mean ± SE. * *P* < 0.05, ** *P* < 0.01. SCr, serum creatinine; Blood urea nitrogen, BUN.

### Urinary exosome NHE3 level in volume depletion induced AKI

The low salt with furosemide injection was recognized as a volume deletion (VD) ^17^AKI model. The raise of SCr (25±1.2 vs. 38±1.8 umol/l, *P*<0.05) and BUN(10.4±1.2 vs. 2.3± 0.4mmol/l, *P*<0.05) meant the AKI model successful (Figure 3(a) and 3(b)). The elevation of urine output after furosemide was in accordance with volume deletion (Figure 3(c)). The pathological injury in VD AKI included the glomerular capillary loop shrinkage and tubular epithelial cell loss (Figure3(e)). We detected the kidney total NHE3 protein by immunoblot, there was no change between control and VD group (Figure 3(f)). Otherwise, the exosome NHE3 from all 30hr urine was increased at VD group(589.3±173 vs. 100.3±36.8, *P*<0.05) (Figure 3(g)). To detect whether it could be as an early biomarker, we split all urine into different periods: 0-8hr,8-24hr and 24–30hr urine. Urine exosomal NHE3 did note elevated at 0-8hr or 8-24hr, though increased at 24-30h(355.5±83.9 vs. 100±22.6,, *P*<0.01) (Figure 3(h)).

**Figure 3.**
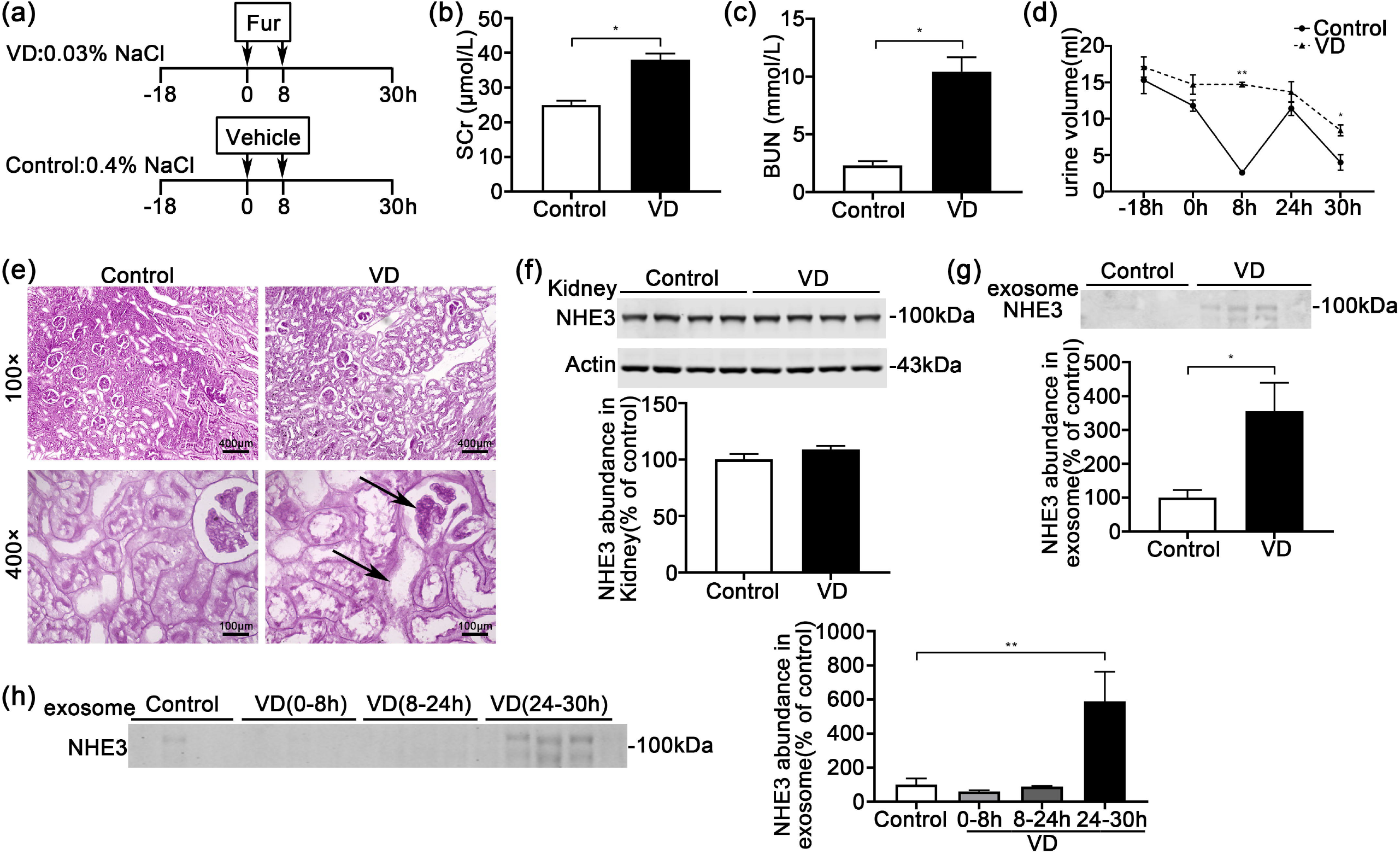
The NHE3 expression in the volume depletion (VD) induced AKI model. (a) The steps of VD-AKI model. The VD group rats were fed food with 0.4% NaCl from −18h to 30h and intraperitoneal injected furosemide (20mg/kg) 2 times at 0h and 8h, while control group were fed with 0.03% NaCl food and injected vehicle at the same time point (n=4/group). (b,c,d) Renal function indicators serum creatinine levels (b), BUN(c), and urine volume (d)were measured. (e) Representative micrographs of histological analysis (PAS staining) of renal tissue from control and VD AKI rats. (f) Immunoblot analyses and quantification of NHE3 protein level in kidney. (g) Urinary exosome NHE3 protein expression between control (all 48hrs urine) and VD-AKI rats(24-30hr). (h) Urinary exosome NHE3 in VD rats in different period: −18-0hr, 0-8hr, 8-24hr and 24–30hr. The data are presented as means ± SE (n =4/group). * *P* < 0.05, ** *P* <0.01vs. vehicle control group. Fur: furosemide; hr: hour.

### Urinary exosome NHE3 excretion in low NaCl with candesartan induced AKI

The SCr (76±10.3 vs. 45.7±2.7 umol/l, *P*<0.05) and BUN(42.2±9.1 vs. 16.7±0.9, *P*<0.05) were elevated at day 7 in low sodium NaCl with candesartan (LS+Can) group (Figure 4(b) and 4(c)). The urine output decreased gradually in both group but no statistic difference at day7 (Figure 4(d)). Histological analysis of renal tissue showed foam cells gathering at tubular epithelial cell, indicated tubular injury (Figure 4(e)). NHE3 protein in kidney tissue(228.1±39.3 vs. 100±24.2, *P*<0.05) was increased in LS+Can group compared with NS+Can group (Figure 4(f)). Urinary exosome NHE3 protein (254.8±24.5 vs. 100± 36.1, *P*<0.01) was markedly increased (Figure 4(g)), and the trend was similar to that in kidney tissue. We detected the urinary exosome NHE3 expression at day 0, 1, 3, 5, 7, found it was incremental, and elevated at day 5(293±7.2 vs.100±22.8, *P*<0.05), which 2 days before the occurrence of SCr increase, still elevated at day 7(702.8±108.1 vs. 100±22.8, *P*<0.01). (Figure 4(h)).

**Figure 4.**
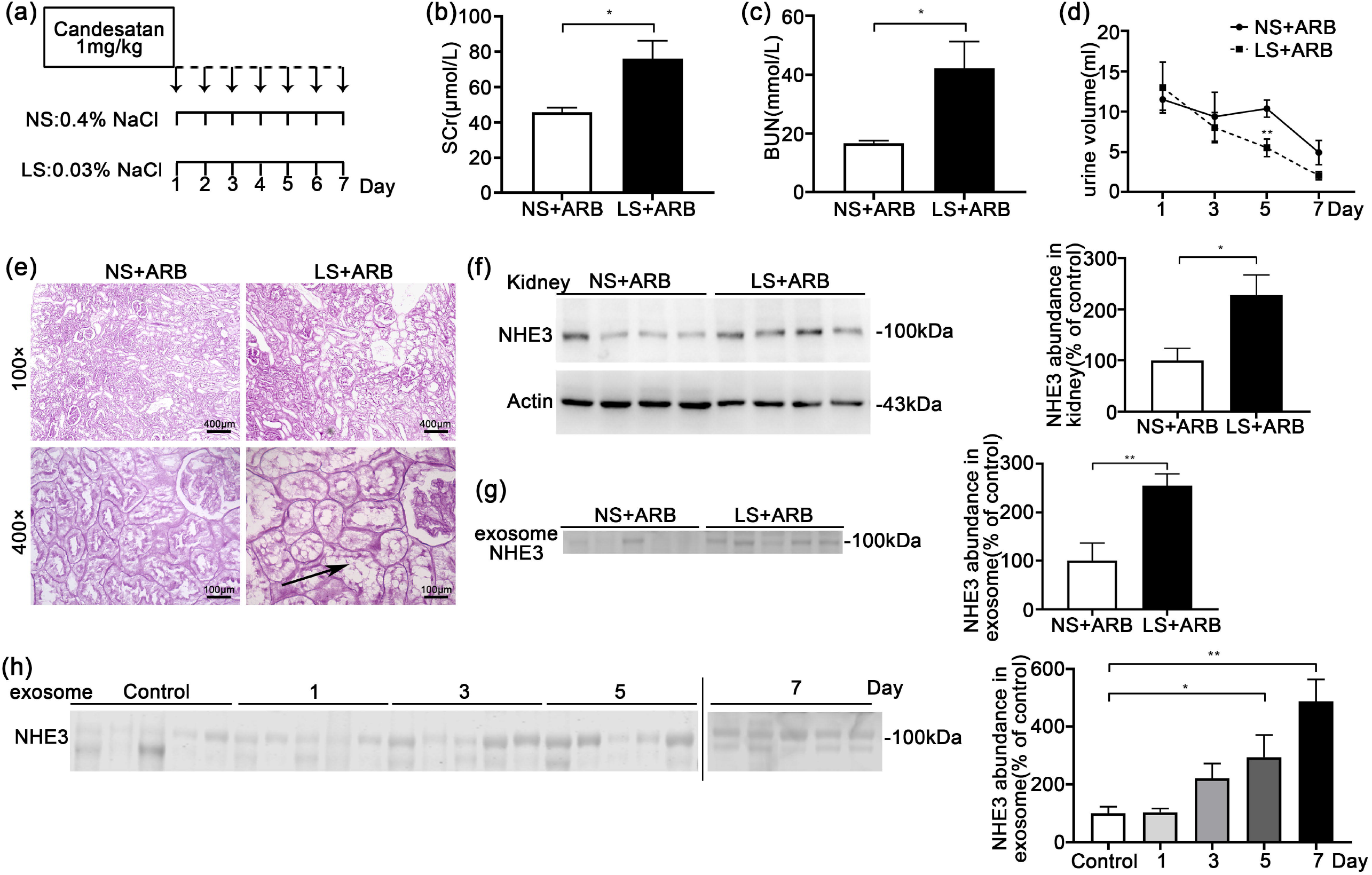
The NHE3 expression in the low NaCl diet with candesartan induced AKI rats. (a) Methods of AKI induced by low NaCl and candesartan. SD rats were intraperitoneally injected with candesartan (1mg/kg/day) for one week, and fed food with low salt with NaCl content at 0.03% (LS+ Can) or normal salt with NaCl content at 0.4% (NS + Can). All rats were sacrificed at the 7th day. (b, c) Changes of serum creatinine, and BUN at day 7 between NS and LS + Can group. (d) The urine output of day1, 3, 5, 7 of the two group rats. (e) Histological analysis (PAS staining) of renal tissue from NS + Can and LS + Can rats. Arrows indicate renal tubular foam cells. (f, g) Immunoblot analyses and quantification of NHE3 protein in kidney f and urine exosomes g at day 7 from NS + Can and LS + Can rats. (h) Urinary exosome NHE3 expression at day 0, 1, 3, 5, 7 from LS + Can induced AKI rats. The data are presented as means ± SE (n =4-5). * *P* < 0.05, ** *P* <0.01.

### Urinary exosome NHE3 level in I/R AKI

The SD rats were bilaterally clamped for 40 min and reperfusion for 24hr. The SCr(270.8± 76.5 vs.22±1.0 umol/l, *P*<0.05) and BUN(34.7±8.2 vs. 4.3±0.2 mmol/l, *P*<0.05) were prominently elevated (Figure 5(a) and 5(b)). The urine volume was declined but no statistic difference (Figure 5(c)). The histomorphological change of renal tissue showed glomerular capillary loop shrinkage, expansion of renal tubules and epithelial cell loss (Figure 5(d)). Both the kidney NHE3(195.2±22.6 vs.100±14.1, *P*<0.05) and urinary exosome NHE3(267±44.4 vs.100±31.1, *P*<0.05) were raised at I/R rats compared with control (Figure 5(e) and 5(f)).

**Figure 5.**
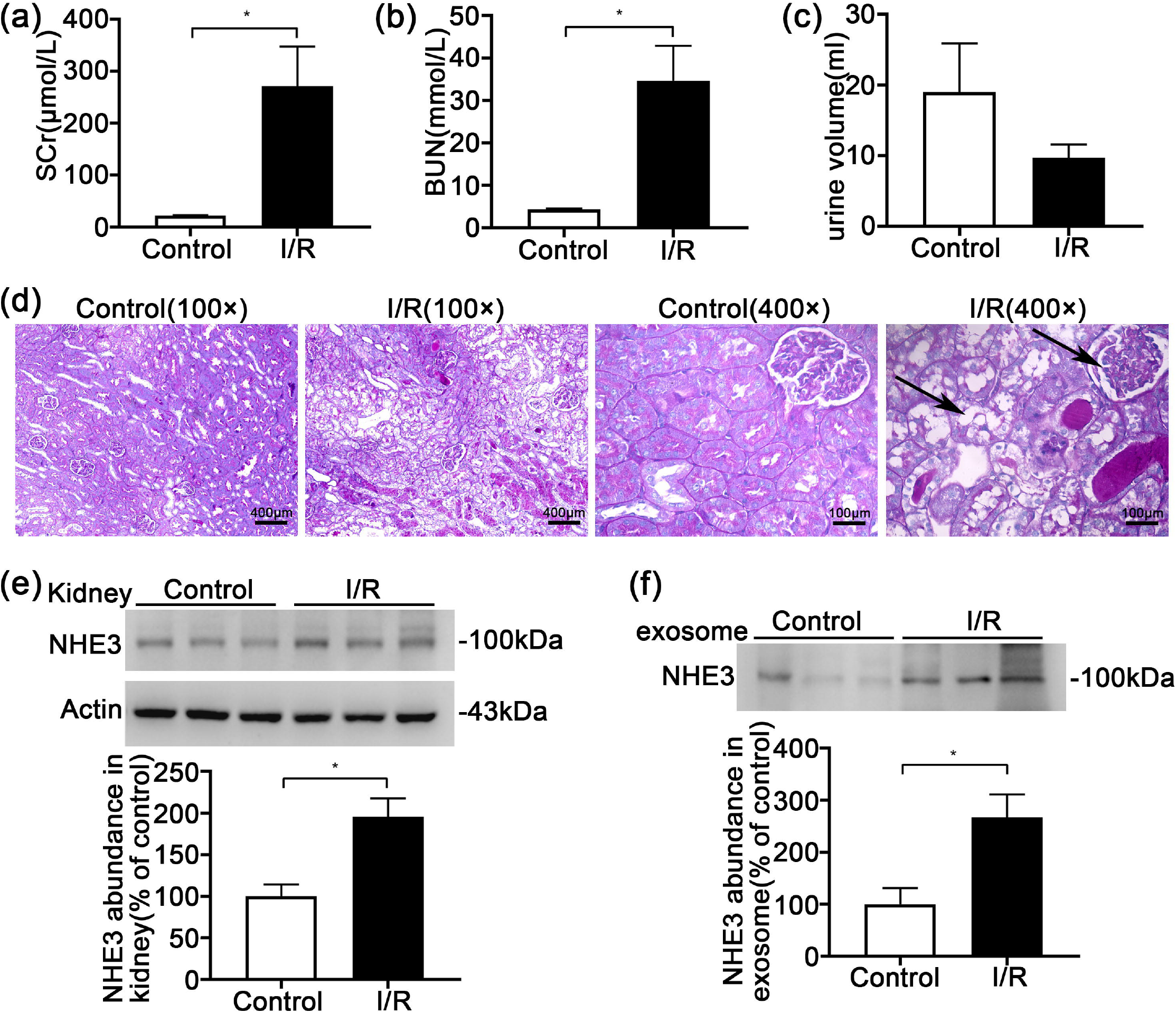
The NHE3 expression in the ischemia/reperfusion(I/R)-induced AKI rats. (a, b, c) Changes of creatinine and urea nitrogen in serum (a, b) and urine output(c) from control or I/R SD rats with bilateral ischemia for 40min, and reperfusion for 24 h. (d) Representative micrographs of histological analysis (PAS staining) of renal tissue from control or I/R rats. (e) Immunoblot analyses and quantification of NHE3 from kidney tissue. (f) Urinary exosome NHE3 protein level between control and I/R rats analyzed by immunoblot. The data are presented as means ± SE (n =3). * *P* < 0.05, ** *P* <0.01 vs. vehicle control group.

### Urinary exosome NHE3 abundance in sepsis associated AKI patients

We chose 6 sepsis associated AKI patients and 6 healthy volunteers. The morning spot urine were collected, and tested urinary exosome NHE3 protein, corrected by the urine creatinine. The general information included age (55±7.2 vs. 58±6.5, P>0.05) and gender (male%,50% vs. 50%, P>0.05) had no difference between AKI and control group. SCr concentrations (258.5±66.9 vs. 68.7±5.4umol/l, *P*<0.05) were significantly increased (Figure 6(a)) and BUN(19.5±3.9 vs. 5.3±0.3mmol/l, *P*<0.01) were elevated (Figure 6(b)). Systolic blood pressure (SBP) (73±2.9 vs. 118.5±3.9 mmHg, *P*<0.01) and diastolic blood pressure (DBP) (41±3.1 vs. 74.8±2.6 mmHg, *P*<0.01) of AKI patients were both lower (Figure 6(c)), and serum C reactive protein (CRP) level was markedly increased (122.7±12.6 vs. 6.6±0.6 mg/L, *P*<0.01) than those in healthy volunteers (Figure 6(d)), that supported the diagnosis of sepsis associated AKI. NHE3 in urinary exosome(1003±89.7 vs. 100±36, *P*<0.01) was also increased significantly compared with the healthy control group (Figure 6(e)).

**Figure 6.**
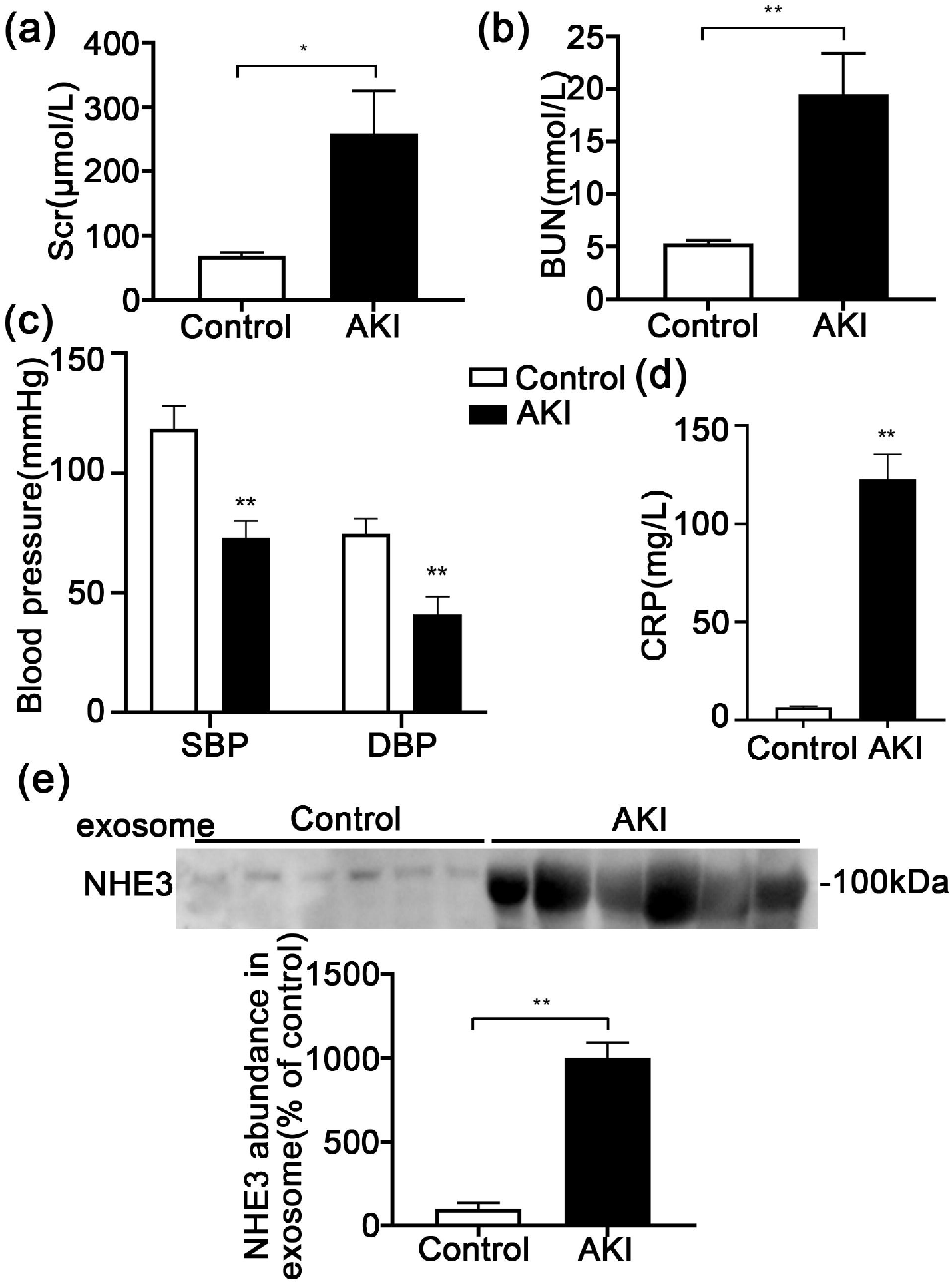
Urinary exosome NHE3 expression in sepsis associated AKI patients. (a, b) The SCr and BUN of sepsis associated AKI patients and healthy volunteers were measured(n=6). (c)Systolic blood pressure and diastolic blood pressure of sepsis associated AKI patients and healthy volunteers. (d) Serum C reactive protein (CRP) level in two group subjects. (e) Immunoblot analyses and quantification of NHE3 from morning urine exosomes. Each value was normalized to the urine creatinine. The data were presented as means ± SE. * *P* <0.05, ** *P* <0.01.

## Discussion

In the present study, we found the urinary exosome NHE3 protein was increased in AKI rats induced by cisplatin, volume depletion, I/R injury and low NaCl with candesartan, and also elevated in sepsis associate AKI patients. Furthermore, urinary exosome NHE3 was detected 1day earlier than SCr in cisplatin-induced AKI rats, and 2 days earlier in low NaCl with candesartan induced AKI. It indicated urinary exosome NHE3 can be used as a noninvasive diagnostic marker of various AKI, and even as an early marker in some type AKI. Exosomes are defined as ∼40 to 160 nm (average ∼100 nm) vesicles in diameter that are secreted when multivesicular bodies fuse with the plasma membrane. ^20^Exosomes are found in all biological fluids and are secreted by all cells. Depending on the cell of origin, exosomes contain many constituents of a cell, including DNA, RNA, lipids, metabolites, and cytosolic and cell-surface proteins. Membrane proteins such as transporters and ion channels are expected to be highly enriched in exosomes^21^.Exosome Detection in biological fluids potentially offers a multicomponent diagnostic readout.^22^ Urinary exosomes are a rich source of biomarkers because they are released from every segment of the nephron^23^. From 2004 Pisitkun Trairak^11^ and the colleagues isolated and identified a series of exosomes protein from healthy human urine, the research about urinary exosomes as biomarkers has risen rapidly.

Exosome Fetuin-A was identified by proteomics as a novel urinary biomarker for detecting AKI^17^, transcription factor ATF3 might be as a renal tubular cell injury biomarker^14^. Urinary exosome CD26^24^ was associated with renal reversal and recovery from AKI. NHE3 was the most abundant sodium transporter in renal tubule, which was localized in the apical membrane and subapical endosomes of renal proximal tubular cells and in the apical membrane of thick ascending limb cells. A few research involved the urinary exosome NHE3^13^, which had some shortage, such as the limited sample size or the complex clinical background. We found NHE3 in urinary exosomes of healthy controls was trace. However, it was increased significantly in various AKI rat model and sepsis-AKI patients. Besides the early diagnosis biomarker, urinary exosome NHE3 might be a recovery marker in cisplatin AKI, which was novel.

The reason of urinary exosome NHE3 increase was worth exploring. In present 4 AKI models, kidney NHE3 protein abundance was elevated in I/R and low NaCl with candesartan, but not in cisplatin or volume depletion induced AKI. Hence, the cause could not be simply attributed to the elevation in renal tissue. Polyuria occurred in volume depletion induced AKI and cisplatin induced AKI, which was associated with a reduction in medullary hypertonicity, due to qualitative changes in urea transporters proteins^19^. Though polyuria might cause the total urinary NHE3 elevation, our exosome loading volume was normalized to the urinary creatinine concentration. Since urinary exosomes were extracelluar microvesicles which were actively secreted by living cells^25^, the increase of urinary exosome NHE3 was considered to be the stress response of proximal tubular epithelial cells to medicine, volume change or others. Besides, NHE3 as the most abundant sodium transporter in renal tubule, which was localized in the apical membrane and subapical endosomes of renal proximal tubular cells and in the apical membrane of thick ascending limb cells, may also appear in the urine as result of incomplete proximal tubule processing in proteinuria states (a form of overflow proteinuria) or released during tubular cell apoptosis.

The cause and pathogenesis of AKI are complex, involving ischemia, sepsis, drug toxicity, and trauma. Cisplatin^26^, ischemia and reperfusion^27^, lipopolysaccharide^28^ and volume depletion^17^ were widely used as AKI animal models. Furthermore, multiple AKI phenotypes exist clinically, and more than one phenotype may exist within the same individual. Low NaCl diet with candesartan was used to study another sodium channel, ENaC, as early as 2005 year^18^, though the kidney function was not noticed. It was a common phenomenon that proteinuria or hypertensive patients often take Angiotensin II receptor blocker with low NaCl intake clinically. We observed the kidney function was damaged at SD rats or spontaneously hypertensive rat, then we established a new AKI model using low NaCl diet with candesartan injection and explored the mechanism. Low NaCl intake with candesartan might promote nitric oxide product and accelerate the hypotension, ultimately induced the AKI (the data was unpublished). In this experiment, we detected the urinary exosome NHE3 in this new AKI model, which was increased 2 days earlier than SCr increase. Urinary exosome NHE3 was raised in multiple AKI models, including the new and complex AKI model, indicating that NHE3 as an AKI marker might be universal.

In the I/R injury AKI rats, the urinary exosome NHE3 protein was detected at 24 hours, as early as the SCr increased. Clamping of renal arteries/vein or pedicle can be performed in two methods-unilaterally or bilaterally. Bilateral clamping of renal arteries, which we took in the experiments, tend to influent the total renal mass and elevate the Scr and BUN levels within 24 h, which are the characteristic features of AKI in a clinical setup^14^. Our original intention was to search earlier markers in I/R AKI rats, but the urine output during 0-12 hours was quite few due to anesthesia and operation, not enough to extract urinary exosomes or test urine creatinine. Therefore, this study did not emphasize early markers in I/R AKI. It still had the practicability and convenience using urine to avoid taking blood frequently.

Due to the limited urine volume in AKI, we did not compare urinary exosome NHE3 with other new biomarkers, such as kidney injury molecule-1, N-acetyl-beta-D-glucosaminidase, neutrophil gelatinase-associated lipocalin, liver fatty acid-binding protein, or the urinary product of tissue inhibitor metalloproteinase and insulin growth factor binding protein-7.

Besides, the mechanism of exosome NHE3 increase in different models of AKI remained to be explore in future. The sample size of AKI patient was relatively small. A larger sample set of AKI patients was needed in the future validation.

In conclusion, urinary exosome NHE3 was elevated in various AKI rats and sepsis associated AKI patients, and increased earlier than SCr in cisplatin induced AKI and low NaCl with candesartan related AKI.Urinary exosome NHE3 may be used as a predictive diagnosis biomarker of AKI in the future.

## AUTHORS’ CONTRIBUTIONS

YTY initiated/designed the study, wrote the manuscript. ZYR and PX completed mouse experiments. ANX did the human part experiment. YTJ performed histopathological staining. YX figured the data according experiments. DXJ conceptualized and designed the entire study. XYW wrote/revised manuscript and is responsible for the integrity of all data included. All authors read and approved the final version of submitted manuscript.

## ETHICAL APPROVAL

The study was approved by the Institutional Review Board and Medical Ethics Committee of Nanjing BenQ Medical Center, The Affiliated BenQ Hospital of Nanjing Medical University (Approval No. 2022-KL005). Written informed consent was obtained from AKI patients and healthy subjects.

